# Community assembly as a basis for tropical forest restoration in a global change scenario

**DOI:** 10.1101/2020.04.04.022400

**Authors:** João Augusto Alves Meira-Neto, Neil Damas de Oliveira-Júnior, Nathália Silva, Ary Teixeira de Oliveira-Filho, Marcelo Leandro Bueno, Vanessa Pontara, Markus Gastauer

**Author notes:** Corresponding author: João Augusto Alves Meira-Neto. **Author contributions:** JAAMN conceived, and designed the research; ATOF conceived the initial database; NDO, NS, MLB, ATOF, and VP built the database; JAAMN, and MG analyzed the data; MG contributed with statistical tools; JAAMN, NDO, NS, MLB, VL, and MG wrote, and edited the manuscript.

## Abstract

Native tropical forests hold high levels of diversity, challenging forest restoration of large areas in a global change scenario. For a site-specific restoration is required the understanding of the main influences ruling the assemblages. We aimed to answer three questions. 1) how do environmental variables influence taxonomic, phylogenetic diversities, and the phylogenetic structure in the of Rio Doce Basin (TFRD)? 2) How do environmental variables, phylogenetic structure and the main types of seed dispersal relate to each other? 3) Which information of the TFRD assemblages can be used for ecological restoration and conservation? We used 78 sites with their checklists to calculate taxonomic, and phylogenetic diversities, phylogenetic structures, and dispersal proportions. Then, we related the diversities of the sites to their bioclimatic variables and built GLM models. Species richness was influenced negatively by water excess duration, by water deficit duration, and positively by maximum temperature, and temperature seasonality. Water regime drives diversity and phylogenetic community structure in the TFRD more than other variables. Annual precipitation and maximum temperature presented the clearer influences on diversity and phylogenetic structure. Zoochory was positively, and anemochory, autochory were negatively related to sesMPD. By choosing the lineages with high fitness for each site, the functioning and the stability of ecosystems should increase. The addition of species with anemochory and autochory increases functional and phylogenetic diversity in areas with extreme water excess or water deficit, important in a global change scenario. A high proportion of zoochory allows the communities to function conserving dispersers, biodiversity, and services.

**Implications for practice:**

- The use of objective methods based on community assembly rules enhances the choice of species, and of phylogenetic lineages better fitted to the bioclimatic profiles of the areas to be restored, improving the functioning and stability of the restored forests.
- The water purification service should be improved through forest restoration as much as possible because ecosystem services and biodiversity conservation are co-benefits of restored forests.
- The inclusion of species with anemochory, and autochory in forest restoration practices should become usual, as they increase functional, and phylogenetic diversities, and as a consequence, the ecosystem stability.
- A large proportion of species with zoochory in restored forests co-benefits taxonomic diversity, phylogenetic diversity, and ecosystem stability.

## Introduction

Environmental degradation is one of the largest threats that are being looked at in the current global change scenario. Large-scale land degradation by agriculture, urbanization and mining are increasing in the tropics (Nazareno & Vitule 2016; Nunes et al. 2019) and require forest restoration activities to reduce the loss of biodiversity and the loss of ecosystem services (Bustamante et al. 2019; Ellison et al. 2017; Filoso et al. 2017; Matos et al. 2020; Poorter et al. 2016). In the Brazilian Rio Doce Basin, the largest river basin (86,715 km^2^) fully inside the Atlantic Tropical Forest Domain, Brazil, a history of land degradation including a recent tailing dam collapse in 2015 (Edwards & Laurance 2015; Meira et al. 2016; Meira-Neto & Neri 2017), accumulated a vegetation deficit of 7,160 km2 to achieve compliance with actual environmental legislation (Pires et al. 2017). Native vegetation of the basin, mainly seasonal tropical forest, holds high levels of diversity (Neves et al. 2020), challenging the restoration of such large areas. Besides choosing the adequate site-specific restoration strategy (Brancalion et al. 2016; Holl & Aide 2011), the selection of locally adapted species is necessary to restitute taxonomic, functional and phylogenetic diversities (Li et al. 2018; Temperton et al. 2005), safeguarding community stability and the performance of ecosystem services (Kang et al. 2018; Pillar et al. 2013), especially in global changing scenarios (Bakker & Wilson 2004; Harris et al. 2006; Wolff et al. 2018). Thus, the understanding of the environmental influences ruling the vegetation assemblages is required to select locally adapted species for planting or seeding (Gastauer et al. 2018).

For deeper understanding of which factors are playing in the assemblage of those Tropical Forests of the Rio Doce Basin (TFRD), rules of community assembly can be assessed by means of tree species composition of forest remnants responding to environmental variables, influencing the taxonomic diversity as well as phylogenetic diversity and structure. This is because (Niño et al. 2014) ecological niches are major drivers of tropical communities largely determined by environmental variables that limit in different ways the species distributions (Götzenberger et al. 2012; Weiher et al. 2011; Weiher & Keddy 1995).

Among the most usable niche determinants in vegetation are the bioclimatic variables (Fick & Hijmans 2017a; Hijmans et al. 2005) which are capable of model spatial distributions of plant species for many different purposes (Austin 2007; Heringer et al. 2019; Phillips et al. 2006; Wüest et al. 2015) allowing predictions in plant communities based on niches (Carnaval & Moritz 2008; Keddy 1992). Niche conservatism and niche convergence, as well as their effects on phylogenetic structures, are useful tools for empirical studies that evaluate environmental filtering and competition as causes of phylogenetic clustering and phylogenetic overdispersion, respectively (Boukili & Chazdon 2017; Losos 2008; Swenson 2009; Wiens et al. 2010). As bioclimatic variables can filter in, and out lineages that have niches conserved or convergent, the phylogenetic structure may varies depending on the phylogenetic signal of selected traits (Blomberg et al. 2003; Lebrija-Trejos et al. 2010).

Besides ecological niches, the main dispersal types are also central in the community assembly since dispersal limitations can play an important rule for forests (Gilbert & Lechowicz 2004; Hardy 2008; Hubbell 2001; Weber et al. 2014). Dispersal types are not only important as drivers of community assembly, but also can inform how these types are aligned with phylogenetic lineages and how these types are conserved or not in the phylogenetic lineages (i.e., phylogenetic signal) (Carrión et al. 2017; Chen et al. 2018).

Therefore, the knowledge about the community assembly of tropical forest communities with high biodiversity is essential for planning of restoration actions, increasing the stability and functioning of the restored and conserved ecosystems (Mori 2016; Oliver et al. 2015; Reich et al. 2012). Since phylogenetic ecology can shed light on the understanding of rules of community assembly and ecological functioning of tropical forests (González-Caro et al. 2014; Oliveira et al. 2014; Paine et al. 2012), we aimed to answer the following questions. 1) How do bioclimatic variables influence taxonomic, and phylogenetic diversities, and the phylogenetic structure in the tropical forests of a river basin (TFRD)? 2) How do environmental variables, phylogenetic structure and the main types of seed dispersal relate to each other in the TFRD assemblages? 3) What of the answers to the previous questions can be used in the ecological restoration of TFRD, as well as in the conservation of biodiversity and ecosystem services?

## Material and Methods

### Study sites/surveys

This study was performed on Tropical Forests of the Rio Doce Basin (TFRD), located between latitudes 17°45’ and 21°15’ S and longitudes 39°30’ and 43°45’W, with more than 86.000 km^2^ occupying parts of Minas Gerais and Espírito Santo states in Brazil (Fig 1). The TFRD area may be influenced by different floras (i.e., Cerrado and Campos Rupestres) on western watersheds, and mountain tops, but Atlantic Tropical Forest flora defines the basin within the domain of Atlantic Forest. We analyzed woody species from subtypes of Atlantic Tropical Forests: seasonal tropical forest and tropical rainforest.

**Figure 1.**
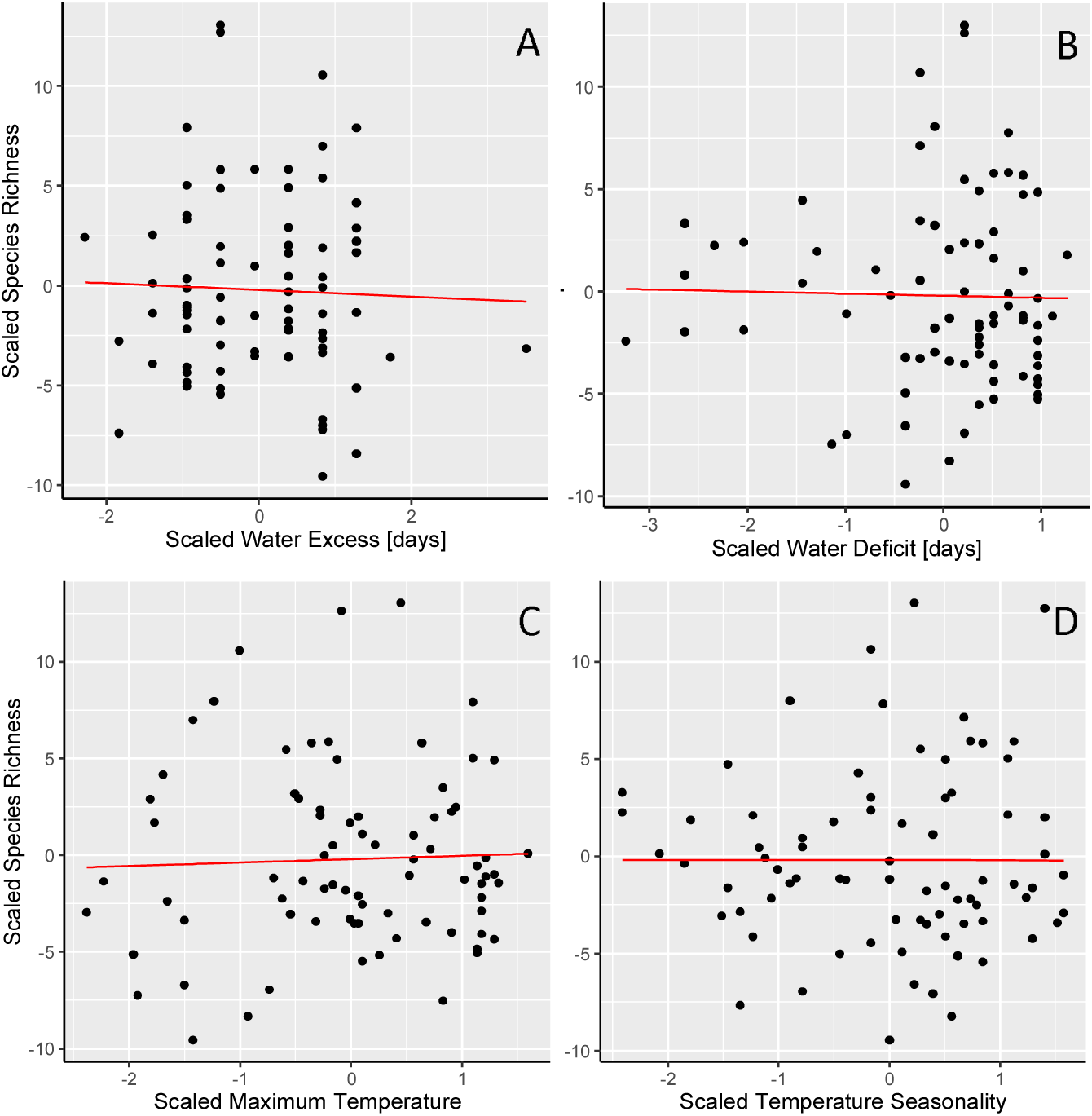
The explanatory A- water excess duration (Water excess [days]), B- water deficit duration (Water deficit [days]), C- maximum temperature of the warmest month (Maximum temperature) and D- temperature seasonality and the response variable species richness of the global GLM model for fragments of the Rio Doce Basin, Brazil. Water excess duration (estimate = -0.108760, P= 4.82e-15), water deficit duration (estimate = - 0.108465, P< 2e-16), maximum temperature of the warmest month (estimate = 0.073362, P= 4.47e-06), temperature seasonality (estimate = 0.065411, P=1.45e-13).

The seasonal tropical forest is defined by a seasonality characterized by a rainy season characteristic of tropical forests followed by a dry season during the winter. As a response to that seasonality, around 20-50% of the trees lose their leaves during the dry period (deciduous). The tropical rain forests are under tropical climate conditions with average temperature of 25°C, and with a very well distributed precipitation along the year, characterized by tall evergreen trees (Oliveira-Filho & Fontes 2000).

The TFRD limits are watersheds of Serra Negra, and Aimorés mountains northwards, the Espinhaço Range on west, the Brigadeiro, and the Caparaó Ranges on southeast, and the ocean in the east. The rainfall regime in the basin is generally divided into rainy season during the summer, and dry season during the winter. The Mean Annual Precipitation varies from 1200 to 1900 mm. The soil of the TFRD are predominantly dystrophic Red-Yellow latosols, deep soils characterized by high aluminium contents, and dystrophic Red Cambisols, a type of shallower soils (Marangon et al. 2013; Nunes et al. 2000; Soil Department of Universidade Federal de Vicosa 2020).

The database has 78 locations of TFRD extracted from NeotropTree, with checklists of tree species (Oliveira-Filho 2014). The checklists are records of papers, monographs, dissertations and thesis as well herbaria records available in the Flora and Fungi Virtual Herbarium (2020). Because the high number of checklists for some sites, the data was compound into one checklist list. A total of 22007 trees of 1944 species within 100 families were analyzed for TFRD (Oliveira-Junior et al. 2020).

### Phylogenetic community structure

We produced a phylogenetic tree with all woody angiosperms recorded in the basin using Phylomatic function in Phylocom 4.2 (Webb & Donoghue 2005) and megatree R20160415.new (Gastauer & Meira-Neto 2017) based on APG IV (The Angiosperm Phylogeny Group 2016). Ferns and gymnosperms were omitted because of their outcome on the phylogenetic metrics they are very distinct and old phylogenetic lineages.

We used the following metrics to explain phylogeny of TFRD in each community: (i) Faith’s phylogenetic Index (PD) (Faith 1992) found by the sum of phylogenetic branches for all species; (ii) Mean Pairwise Distance (MPD), the mean phylogenetic distance among species; (iii) Mean Nearest Taxon Distance (MNTD) (Webb 2000; Webb et al. 2002), the mean phylogenetic distance between the closest related species. Also, we calculated the standardized effect size (ses) of PD, MPD and MNTD to eliminate the richness effect based on a null model (Swenson 2014). Negative ses show phylogenetic clustering and positive ses show phylogenetic overdispersion.

We used evolutionary distinctiveness scores (ED) as species exclusivity (Isaac et al. 2007). If a species does not have close relatives in a community phylogeny, that species is considered unique and ED scores high (Edwards et al. 2017).

### Environmental variables

For each localization of the surveys, we gathered elevation (United States Geological Survey 2019) and extracted the 14 bioclimatic variables from WorldClim: Annual Mean Temperature, Isothermality, Temperature Seasonality, Maximum Temperature of the Warmest Month, Minimum Temperature of Coldest Month, Mean Temperature of Wettest Quarter, Annual Precipitation, Precipitation of Wettest Month, Precipitation of Driest Month and Precipitation Seasonality (Fick & Hijmans 2017b). Additionally, we used WaterExcDur, WaterExcSev, WaterDefDur and WaterDefSev from diagrams of Walter (Oliveira-Filho 2014; Walter 1986; Walter & Lieth 1967) for modelling. All bioclimatic variables are derived from the monthly or daily temperature and rainfall values in order to produce ecologically important variables used for niche modelling with high consistency (García-Callejas & Araújo 2016; Morales-Barbero & Vega-Álvarez 2019). The applied bioclimatic variables signify annual means (Booth 2016; Kendal et al. 2018), seasonality (Hereford et al. 2017; Tonkin et al. 2017) and severe environmental factors (e.g., duration and severity of water excess and deficits) that influence species distribution and performance (e.g., Amissah et al., 2018).

### Phylogenetic signal of dispersal types

Phylogenetic signal of dispersal types means the trend of species with zoochory, anemochory, and autochory to be more related than by chance or to be less related than by chance. If the signal shows the pool of species with a dispersal type more related between each other than null models, the phylogenetic signal is clustered and means that the dispersal type has been predominantly conserved within the phylogenetic lineages under niche conservatism. The opposite, when a signal shows the species less related than calculated by null models, the phylogenetic signal is overdispersed and means that the dispersal type is predominantly convergent (homoplasy). In order to test if dispersal types presented phylogenetic signal, the NRI among all species of the same dispersal type (DispersalNRI) was calculated. The same was calculated for NTI (DispersalNTI). This calculation was done reordering species labels 999 times through phylogeny. DispersalNRI values significantly greater than zero (more than 1.96 sd, and more than -1.96 sd) indicate phylogenetic signal. Phylogenetic signal of dispersal type was considered significant when the observed MPD or MNTD occurred in the lowest 2.5% of the null models (significance of 0.05) (Gastauer et al. 2017).

### Modelling

To outline the environmental predictor variables that best describe species richness and phylogenetic structure of sampled tree communities, we build for each response variable a global general linearized model containing all 15 predictor variables.

We scaled the explaining variables. Then, we used the dredge function of the MuMIn package (Bartón 2018) to model-select all possible combinations of five or less uncorrelated predictive variables (r <0.6, https://github.com/rojaff/dredge_mc) that most parsimoniously explained our data. We considered models with five or less predictive variables to avoid overfitting. We used the Akaike Information Criterion (AIC) for selection of the best models (Symonds & Moussalli 2011). All models with ΔAIC less than 2 were considered equally parsimonious. When more than one model was selected, we calculated model-averaged parameters and unconditioned standard errors weighted by the likelihood ratio using the ‘model.avg’ function of the MuMIn package. The function performs significance tests for each predictor estimated in this average model. The resulting global models were confirmed by plotting residuals versus adjusted values as well as residuals versus predicts.

For modelling percentages of zoochorous, anemochorous and autochorous species within surveys, we build global models containing the same 15 environmental predictor variables plus species richness, ses.MPD, ses.MNTD and ses.PD. For model selection, we proceed as described above.

## Results

Modelling species richness, only one global model containing five environmental variables was selected, with four significant trends (Table S1), indicating that richness is influenced negatively by water excess duration and by water deficit duration meanwhile is positively influenced by the Maximum temperature of the warmest month as well as is positively influenced by seasonal temperature (Figure 1).

For further response variables, more than one model was selected (Table S1). 14 models containing the all variables except Precipitation during Wet Period, and Minimum Temperature explain sesPD. However, only two trends were significant: annual precipitation reduces sesPD, and water excess duration increases sesPD (Table S2, Figure 2).

**Figure 2.**
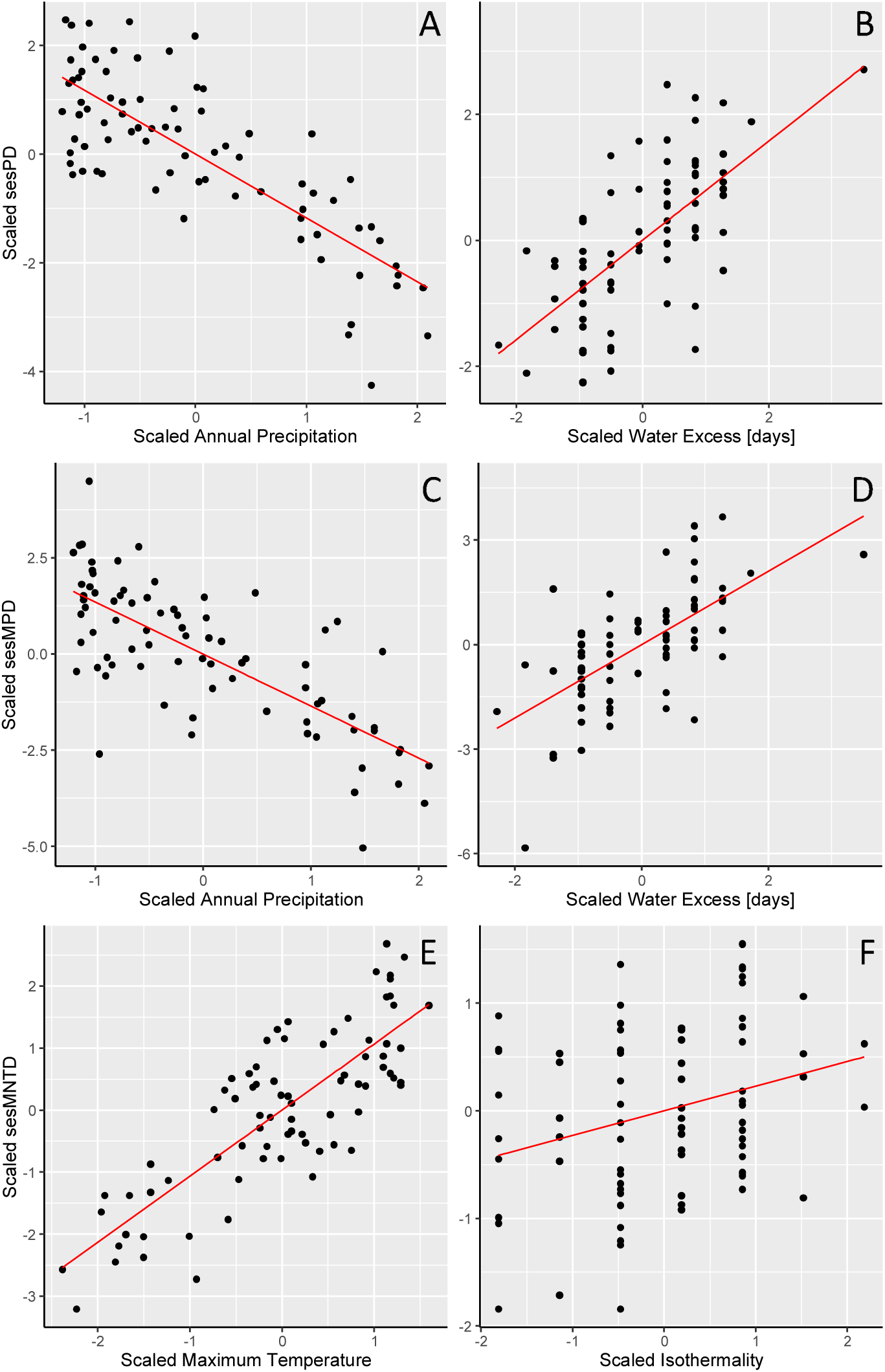
A, and B - The explanatory variables mean annual precipitation (Annual precipitation) and water excess duration (Water excess [days]), and the response variable standard size effect of phylogenetic diversity (sesPD) of the global GLM model for fragments of the Rio Doce Basin, Brazil. Annual precipitation (estimate = - 1.173858, P=0.003); Water excess (estimate = 0.786778, P=0.00332). C, and D- The explanatory variables mean annual precipitation (Annual precipitation) and minimum temperature of the coldest month (Minimum Temperature) and the response variable standard size effect of mean phylogenetic distance (sesMPD) of the global GLM model for fragments of the Rio Doce Basin, Brazil. Annual precipitation (estimate = -1.353912, P=0.00357); Water excess (estimate = 1.052365, P=0.00368). E, and F- The explanatory variables maximum temperature of the warmth month, and the isothermality, the response variable of sesMNTD of the global GLM model for fragments of the Rio Doce Basin, Brazil. Maximum temperature (estimate = 1.06510, P=0.00389), isothermality (estimate = 0.22868, P=0.04736).

For ses.MPD, six models were selected containing 10 out of the 15 predictor variables (Tables S1, S2). In the averaged model, only two trends were significant: annual precipitation reduces and water excess duration increases sesMPD (Figure 2).

For ses.MNTD, the two significant trends were: increasing maximum temperature of the warmest month, and increasing isothermality increase sesMNTD (Table S2, Figure 2).

The percentage of zoochory species increases with increasing sesMPD, with water excess duration, with water deficit severity, and decreases with water excess severity (Figure 3).

**Figure 3.**
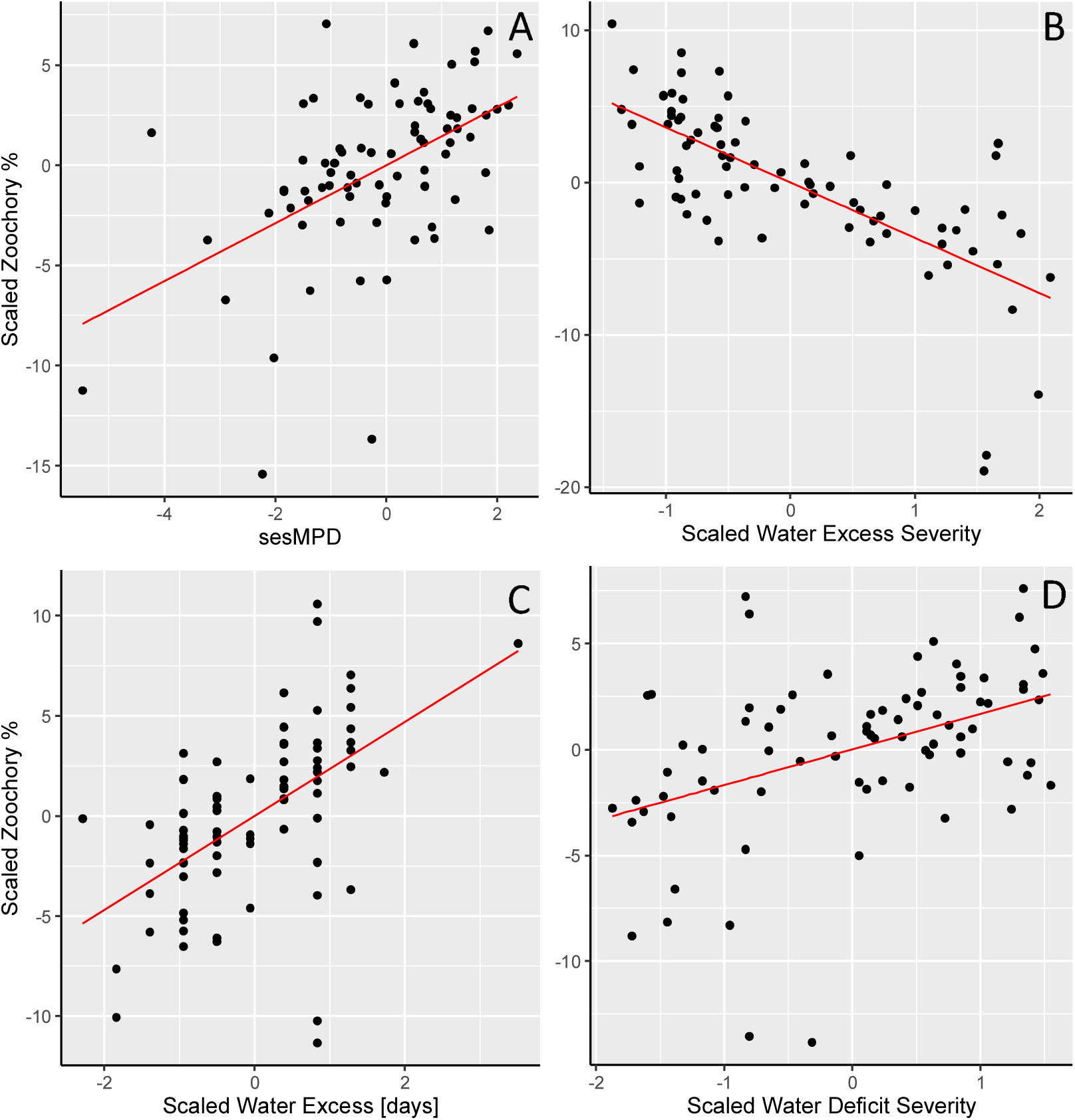
The explanatory variables standard size effect of A- mean phylogenetic distance (sesMPD), B- water excess severity, C- water excess duration (Water excess [days]), D-water deficit severity, and the response variable percentage of species with zoochory (Zoochory %) of the global GLM model for fragments of the Rio Doce Basin, Brazil. SesMPD (estimate = 1.4483, P= 1.56e-05), water excess severity (estimate = -3.6250, P= 1.84e-05), water excess duration (estimate = 2.3493, P= 0.0144), water deficit severity (estimate = 1.6785, P=0.01964).

The percentage of species with anemochory decreases with increasing sesMPD and increases with increasing water excess severity (Table S3, Figure 4).

**Figure 4.**
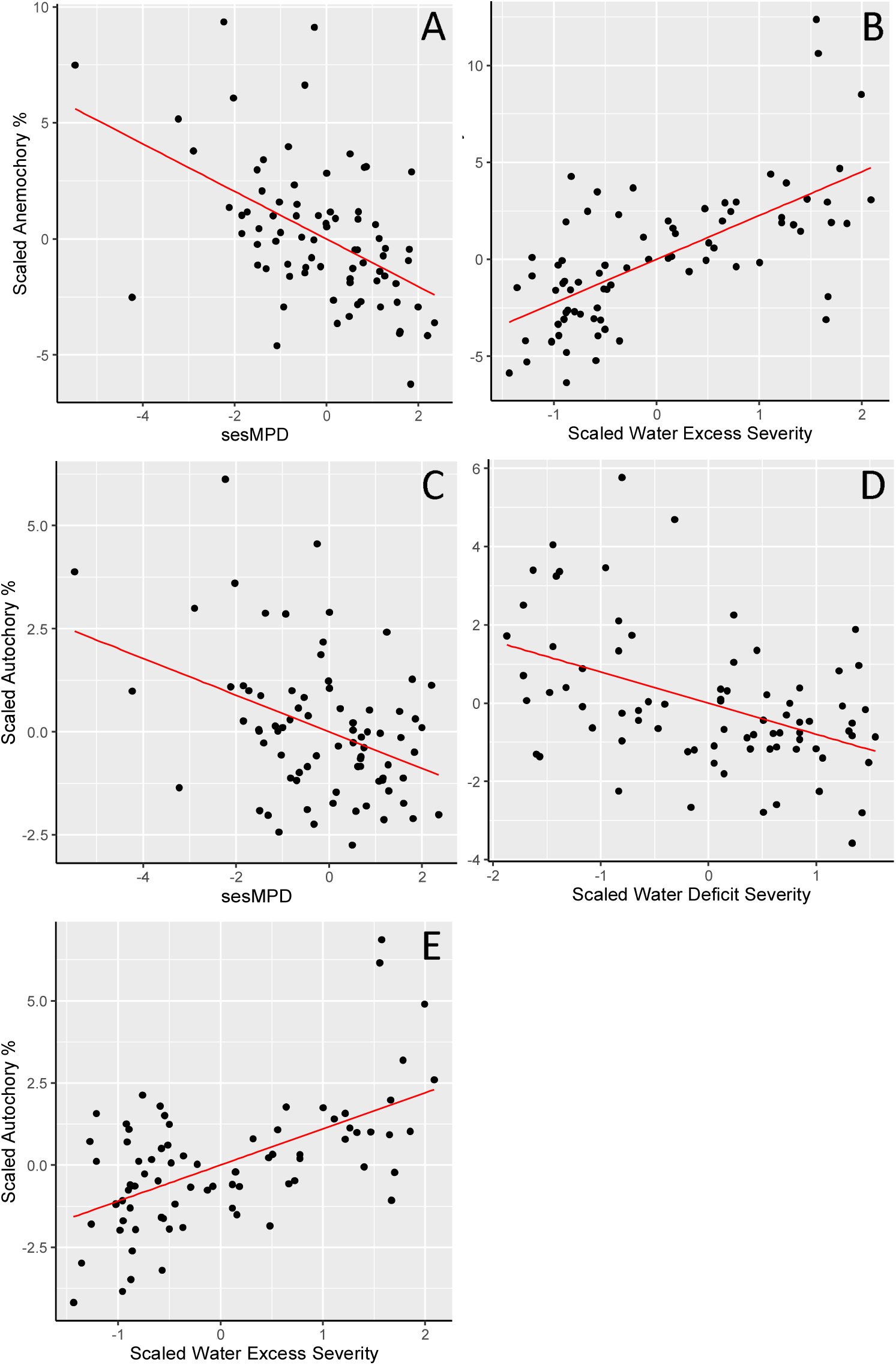
A, and B- The explanatory variables standard size effect of mean phylogenetic distance (sesMPD), water excess severity and the response variable percentage of species with anemochory (Anemochory %) of the global GLM model for fragments of the Rio Doce Basin, Brazil. SesMPD (estimate = -1.02441, P= 4.82e-05), water excess severity (estimate = 2.26164, P= 0.00181). C, D, and E- The explanatory variables standard size effect of mean phylogenetic distance (sesMPD), water deficit severity, and water excess severity, and the response variable percentage of species with autochory (Autochory %) of the global GLM model for fragments of the Rio Doce Basin, Brazil. SesMPD (estimate = -0.4452, P=0.00186), water deficit severity (estimate = - 0.7936, P=0.00935), water excess severity (estimates = 1.1008, P=0.00188).

The percentage of autochory increases as sesMPD decreases, as water deficit severity decreases, and is positively related to the water excess severity (Table S3, Figure 4).

Anemochory, and autochory presented DispersalNRI clustered phylogenetically showing that these dispersal types are conserved within phylogenetic lineages throughout the phylogenetic tree (Table 1). Zoochory, and anemochory preented clustered DispersalNTI, showing that hese dispersal types are conserved within phylogenetic lineages towards the tips of the phylogenetic branches (Table 1).

**Table 1.** Phylogenetic signal of dispersal types as DispersalNRI (MPD), and DispersalNTI (MNTD) for anemochory, autochory, and zoochory. Dispersal types: AN - anemochory, AU - autochory, and ZO - zoochory. Calculated by Phylocom with randomization method 2, by mean of 999 runs. Ntaxa, number of species; MPD, mean phylogenetic distance calculated for species by actual data; MPD.rnd, mean phylogenetic distance calculated by randomization; NRI, Net Relatedness Index; MPD.rank.Low, how many times the MPD was lower than MPD.rnd; MPD.rank.Hi, how many times the MPD was higher than MPD.rnd; MNTD, mean nearest taxon distance calculated for species by actual data; MNTD.rnd, mean nearest taxon distance calculated by randomization; NTI, Nearest Taxon Index; MNTD.rank.Low, how many times the MNTD was lower than MNTD.rnd; MNTD.rank.Hi, how many times the MNTD was higher than MNTD.rnd; runs, number of randomizations. *p<0.05, **p<0.01, ***p<0.001.

| Type | ntaxa | MPD | MPD.rnd | MPD.sd | NRI | MPD.rankLow | MPD.rankHi | runs |
| --- | --- | --- | --- | --- | --- | --- | --- | --- |
| AN | 330 | 209.0239 | 220.9857 | 1.4227 | 8.4076*** | 999 | 0 | 999 |
| AU | 161 | 191.2269 | 221.1116 | 2.0766 | 14.3909*** | 999 | 0 | 999 |
| ZO | 1432 | 221.3495 | 221.1048 | 0.3678 | -0.6653 | 247 | 752 | 999 |
| Type | ntaxa | MNTD | MNTD.rnd | MNTD.sd | NTI | MNTD.rankLo | MNTD.rankHi | runs |
| AN | 330 | 56.0852 | 61.3007 | 1.8922 | 2.7563** | 996 | 3 | 999 |
| AU | 161 | 67.5397 | 71.9932 | 3.4141 | 1.3044 | 899 | 100 | 999 |
| ZO | 1432 | 44.6573 | 48.8249 | 0.3891 | 10.71*** | 999 | 0 | 999 |

## Discussion

Water regime drives species richness and phylogenetic community structure in the TFRD more than other types of environmental variables. At least one among annual precipitation, water excess duration, water excess severity, and water deficit severity were significant in all averaged models, except for the model for sesMNTD. Species richness, sesPD, sesMPD and percentage of zoochory were affected by water excess duration. All dispersal types were explained partially by water excess severity. Temperature regime was the second mst important important regime. Maximum temperature of the warmest month and isothermality explained sesMNTD. Maximum temperature of the warmest month, and temperature seasonality explained partially species richness. Among the variables, water excess duration, annual precipitation, maximum temperature and isothermality influenced the phylogenetic diversity and phylogenetic structure. The effects of temperature on richness, and on sesMNTD suggest environmental filtering effects causing phylogenetic clustering driven by decreasing maximum temperature of the warmest month and decreasing isothermality (i.e., increasing temperature variation). Therefore, the lower temperature and the less temperature variation in different seasons, the more environmental filtering.

The percentage of species with zoochory, anemochory or autochory were related to phylogenetic structures (i.e., sesMPD) in different ways. Increasing zoochory increases phylogenetic distances between species (i.e., increasing sesMPD) meanwhile increasing anemochory, and autochory decrease phylogenetic distances between species (i.e., decreasing sesMPD). The opposite relation of zoochory with sesMPD was not because zoochory is convergent, but because the less zoochory, the more abiotic dispersion (i.e., anemochory + autochory) that is clustered for MPD (i.e., DispersalNRI more related than by chance), suggesting anemochory, and autochory as homologies in the TFRD metacommunity. Only zoochory decreased with increasing water excess severity, and appears to be the only dispersal type benefited with increasing water surplus. Water excess severity was positively related to anemochory, and autochory, and might be interpreted as stresses influencing positively these dispersal types. Water deficit severity influenced negatively autochory.

The answer of the first question shows how environmental variables influence taxonomic, and phylogenetic diversities as well as phylogenetic structure in the tropical forests of that large river basin (TFRD). Seven environmental variables were found to influence the taxonomic diversity, phylogenetic diversity and phylogenetic structure in averaged models: water excess duration, water excess severity, water deficit duration, water excess severity, maximum temperature, temperature seasonality, and isothermality. As water excesses and water deficits increase in number of days, the species richness decreases. As the maximum temperature and the temperature seasonality decrease, the species richness decrease. Thus, more days of water excesses or more days of water deficits in TFRD are associated with loss of species richness as well as lower maximum temperatures and lower temperature variation between seasons. Therefore, the found global model shows that increasing the number of days with excess or deficit of water, decreasing temperatures, and decreasing temperature variation, decrease species richness.

Still answering the first question, as annual precipitation decreases and there are more days of water excess, sesPD and sesMPD increase; as maximum temperature and isothermality increase, sesMNTD increases. Thus, the lower annual precipitation, the more water excess days, the higher temperature variation, and the higher maximum temperatures in TFRD, the greater the phylogenetic diversity and phylogenetic distances between species. However, as the more days with water excess (i.e., days with higher precipitation than evapotranspiration) cause higher sesPD and sesMPD, it is possibly because benefits the zoochory that is positively-related to mean phylogenetic distances (see zoochory discussion below). Therefore, higher precipitation promotes decreasing sesPDs and sesMPDs (i.e., phylogenetic clustering) in the TFRD similarly to effects of environmental filters found in tropical vegetation (Gastauer & Meira-Neto 2013; Miazaki et al. 2015), but does not cause species richness loss. Phylogenetic effects in tropical vegetation that can be explained by plant-plant interactions, density-dependence effects (Cadotte & Tucker 2017; Carrión et al. 2017; Kraft et al. 2015; Meira-Neto et al. 2018; Paine et al. 2012), and environmental filtering that has been reported as cause of phylogenetic effects in many different groups of species, especially plants. For instance, environmental filtering promotes phylogenetic effects in protists, in diatoms (Keck & Kahlert 2019; Leibold et al. 2010; Singer et al. 2018), and likely causes phylogenetic clustering in plant communities of tropical forests when associated with species richness decreasing (Gastauer & Meira-Neto 2013; Parmentier et al. 2014). Therefore, the increasing annual precipitation in the TFRD that shortens phylogenetic distances might be an environmental filter effect because of excessive water. High precipitation may cause temporarily flooding stresses (Bueno et al., 2014; Pontara et al., 2016), and, consequently, may filter out lineages without conserved traits of flooding tolerance, but species richness does not decreased with the increasing annual precipitation. Alternatively, increasing annual precipitation would promote competition, and/or density dependent effects. For instance, high precipitation might cause competition increasing, and higher density-dependent effects by increased water resources and increased plant growing (see Tilman 1988). However, the forests with largest trees in TFRD, which would indicate stronger competition, are in low altitude sites with lower precipitation (Camargos et al. 2008; Lopes et al. 2002; Souza et al. 2013, 2012). The only clear result is that as precipitation increases, PD, and MPD decrease as an effect of increasing percentages of species with abiotic dispersion (see dispersal discussion below). Therefore, environmental filtering, competition and density dependent effects do not explain the PD and the MPD variation in TFRD. On the other hand, temperature regime can explain environmental filtering because species richness and sesMNTD decrease as maximum temperature decreases, indicating that the lower the maximum temperature, the closer the relatives in the TFRD communities and the lower the species density. High and low temperatures have been reported as environmental filters in forests, but high temperatures are rather filters in tropical dry forests, and low temperatures are rather filters associated with high altitudes or latitudes in temperate forests (Qian 2018; Qian et al. 2014). Therefore, our results suggest that the seasonal TFRD are environmentally filtered by low temperatures that cause decreasing species richness, and decreasing phylogenetic distance between species of their communities by filtering in species with conserved traits within phylogenetic lineages.

The second question was how the dispersal types relate with environmental variables and with phylogenetic structure in the TFRD assemblages. The percentage of species with zoochory, anemochory and autochory was significantly related to sesMPD and not to sesMNTD or sesPD, which means that the main association of dispersal types with the communities’ phylogenies is throughout the entire phylogenetic tree (i.e., sesMPD). As the DispersalNRI is clustered for abiotic dispersal types, the clustered sesMPD reinforce that anemochory and autochory are conserved in phylogenetic branches originated in old nodes congruently with the decreasing sesMPD as anemochory and autochory. As a consequence, the increasing proportion of zoochory is not caused by its preeminence as homoplasy in the TFRD metacommunity, but because abiotic dispersal decreases. Thus, increasing anemochory and autochory reduce the sesMPD distances between species congruently with their clustered DispersalNRI in the TFRD metacommunity.

Zoochory is phylogenetic clustered towards the tip of phylogenetic tree according to its DispersalMNTD. That means the species with zoochory are predominantly clustered in lineages with conserved zoochory diversified from recent nodes. Despite zoochory is clustered towards the tip of phylogenetic tree, zoochory is not clustered throughout the entire phylogenetic tree. Anemochory is also clustered clustered according to its DispersalMNTD. Autochory is not clustered towards the tip of phylogenetic tree in TFRD, suggesting lack of recent diversification of lineages with conserved autochory.

The dispersal types responded to water regime. The dominant dispersal type in TFRD is zoochory that varies from 50% up to more than 95% of the tree species in evergreen and seasonal Atlantic Tropical Forests (Tabarelli & Peres 2002). In the TFRD, the minimum zoochory percentage found in a forest was 53% and the maximum was 81% (data not shown). As dominant dispersal type, zoochory responded positively to water excess duration, and water deficit severity meanwhile anemochory and autochory were not influenced by those explaining variables. However, zoochory responded in an opposite way from the two abiotic dispersal types to water excess severity. Zoochory also responded in an opposite way from autochory to water deficit severity. Thus, zoochory seems to be impaired meanwhile anemochory and autochory seem to be benefitted in sites with extreme severity of water excess or water deficit.

As anemochory and autochory are clustered in some phylogenetic branches in the TFRD metacommunity, the presence of these dispersal types in restored TFRD will allow those species to improve the functional and phylogenetic diversity of restored forests by representing entire phylogenetic branches, especially in those TFRD with extreme water regimes (i.e., high severity of water excess, and high severity of water deficit). Therefore, the relation between severity of water excess, and severity of water deficit with the two abiotic dispersal types deserves attention since extreme water regimes are increasing according to forecasted climate scenarios in the most of the Atlantic Forest distribution (IPCC 2013). As a consequence, the proportion of anemochory and autochory may increase in the TFRD during the next decades in a global change scenario.

Answering the question about what information from the TFRD community assembly can be used in ecological restoration, biodiversity conservation and conservation of ecosystem services, the results showed that the TFRD community assembly is central to large-scale forest restoration planned in the basin, including changing the effects of the Mariana disaster, a tailing dam that collapsed, and that compromised the service of the water purification in the basin. 1) The environmental filtering promoted by low temperatures (i.e., low maximum temperatures) drops the taxonomic diversity shortening the phylogenetic distance between species, predominantly filtering in species of some recently diversified lineages. Temperatures in TFRD are also related with altitude (data not shown). Thus, the use of some lineages better fitted for TFRD restoration in sites with low maximum temperatures (i.e., high altitudes inside the basin) should enhance the performance of restored forests. 2) Possibly the most affected ecosystem service by forests degradation, agriculture (Cabel et al. 1982), urbanization (Zhao et al. 2013), and environmental disasters (Lambertz & Dergam 2015; Meira et al. 2016; Meira-Neto & Neri 2017) is water purification that should be restored in all regions of the basin by means of forest restoration. We understand that tree species with good fitness to the bioclimatic profiles of restored sites will enhance the ecosystem service of water purification, especially in areas with low maximum temperature, and high annual precipitation, improving the functioning and the stability of restored forests (Cadotte et al. 2012, 2009; Isbell et al. 2015; Brancalion et al. 2016). 3) In a global change scenario that predicts increasing climate variance for TFRD (IPCC 2013), the inclusion of species with anemochory and autochory in restoration practices increases functional and phylogenetic diversity when associated with a large proportion of species with zoochory, especially in areas with extreme water excess or extreme water deficit. 4) A large proportion of species with zoochory in restored forests is mandatory because it assembles communities with high taxonomic, and phylogenetic diversity, conserving dispersal services (Jansen et al. 2014; Tabarelli & Peres 2002), conserving dispersers, and conserving the threatened ecological interactions of tropical forests (Janzen 1986).

## Acknowledgements

The authors thank the Botany Graduate Program (NS, VP) and the Ecology Graduate Program (MLB) for their support. The funding was provided by FAPEMIG (PPM-00584-16, APQ-01309-16), CAPES (PROAP fund and scholarships for NDO, VP, and MLB), and CNPq (NS scholarship, and 446698/2014-8). JAAMN holds a CNPq productivity fellowship (307591/2016-6).

