## Supplementary material for "Community assembly as a basis for tropical forest restoration in a global change scenario"

**Running head:** Community assembly as basis for restoration in a global change

**Authors and adresses:**

João Augusto Alves Meira-Neto^1,2^, Neil Damas de Oliveira-Júnior^1,2^, Nathália Silva^1,2^, Ary Teixeira de Oliveira-Filho^3^, Marcelo Leandro Bueno^1,2,4^, Vanessa Pontara^1,2,4^, Markus Gastauer^1,2,5^

^1^ Universidade Federal de Viçosa, Laboratory of Ecology and Evolution of Plants – LEEP, Viçosa, Minas Gerais, Brazil, 36570-900.

^2^ Universidade Federal de Viçosa, Botany Graduate Program, Viçosa, Minas Gerais, Brazil, 36570-900.

^3^ Universidade Federal de Minas Gerais, Botany Department of Universidade Federal de Minas Gerais, Belo Horizonte, Minas Gerais, Brazil, 31270-901.

^4^ Laboratório de Evolução e Macroecologia, Universidade Estadual de Mato Grosso do Sul, BR 163, Mundo Novo, Mato Grosso do Sul, Brazil, 79980-000.

^5^ Instituto Tecnológico Vale, Rua Boaventura da Silva, 955, Belém, Pará, Brazil, 66055-090.

Table S1. Selected models for predicting the species density, phylogenetic structure and diversity for forest remnants from the Rio Doce Basin

| Response variable | Model parameters | df | logLike | AIC | delta | Weight |
| --- | --- | --- | --- | --- | --- | --- |
| Species richness | Maximum temperature + Water Excess Severity + Temperature Seasonality + Water Deficit Duration + Water Excess Duration | 6 | -1107.82 | 2227.6 | 0.00 | 0.998 |
| ses.PD | Annual Precipitation + Precipitation Seasonality + Minimum Temperature + Water Excess Duration + Water Excess Severity | 7 | -105.238 | 224.5 | 0.00 | 0.067 |
|  | Annual Precipitation + Minimum Temperature + Water Excess Duration + Water Excess Severity + Precipitation Dry Period | 7 | -105.528 | 225.1 | 0.58 | 0.050 |
|  | Annual Precipitation + Water Excess Duration + Water Excess Severity + Mean Annual Temerpature | 6 | -106.548 | 225.1 | 0.62 | 0.049 |
|  | Annual Precipitation + Water Excess Duration + Water Excess Severity + Mean Annual Temperature + Isothermality | 7 | -105.577 | 225.2 | 0.68 | 0.048 |
|  | Annual Precipitation + Water Excess Duration + Mean Annual Temperature + Water Deficit Duration | 6 | -106.723 | 225.4 | 0.97 | 0.041 |
|  | Annual Precipitation + Minimum Temperature + Water Excess Duration + Water Excess Severity | 6 | -106.738 | 225.5 | 1.00 | 0.041 |
|  | Annual Precipitation + Water Excess Duration + Mean Annual Temperature + Isothermality + Water Deficit Duration | 7 | -105.751 | 225.5 | 1.03 | 0.040 |
|  | Annual Precipitation + Precipitation Seasonality + Water Excess Duration + Mean Annual Temperature + Isothermality | 7 | -105.785 | 225.6 | 1.10 | 0.039 |
|  | Annual Precipitation + Water Excess Duration + Water Deficit Duration + Elevation + Temperature Seasonality | 7 | -105.887 | 225.8 | 1.30 | 0.035 |
|  | Annual Precipitation + Precipitation Seasonality + Water Excess Duration + Mean Annual Temperature | 6 | -106.986 | 226.0 | 1.50 | 0.032 |
|  | Annual Precipitation + Minimum Temperature + Water Excess Duration + Water Excess Severity + Isothermality | 7 | -106.068 | 226.1 | 1.66 | 0.029 |
|  | Annual Precipitation + Water Excess Duration + Water Excess Severity + Water Deficit Duration Maximum Temperature | 7 | -106.107 | 226.2 | 1.74 | 0.028 |
|  | Annual Precipitation + Water Excess Duration + Water Excess Severity + Mean Annual Temperature + Water Deficit Duration | 7 | -106.178 | 226.4 | 1.88 | 0.026 |
|  | Annual Precipitation + Water Excess Duration + Water Excess Severity + Maximum Temperature + Water Deficit Severity | 7 | -106.189 | 226.4 | 1.90 | 0.026 |
| ses.MPD | Elevation + Annual Precipitation + Temperature Seasonality + Water Excess Duration | 6 | -130.731 | 273.5 | 0.00 | 0.079 |
|  | Elevation + Annual Precipitation + Temperature Seasonality + Water Excess Duration + Water Deficit Duration | 7 | -130.462 | 274.9 | 1.46 | 0.038 |
|  | Annual Precipitation + Water Excess Duration + Precipitation Dry Period + Mean Annual Temperature | 6 | -131.591 | 275.2 | 1.72 | 0.033 |
|  | Annual Precipitation + Water Excess Duration + Minimum Temperature + Water Excess Severity | 6 | -131.619 | 275.2 | 1.78 | 0.032 |
|  | Elevation + Annual Precipitation + Temperature Seasonality + Water Excess Duration + Water Excess Severity | 7 | -130.667 | 275.3 | 1.87 | 0.031 |
|  | Elevation + Annual Precipitation + Temperature Seasonality + Water Excess Duration + Water Deficit Severity | 7 | -130.730 | 275.5 | 2.00 | 0.029 |
| ses.MNTD | Isothermality + Precipitation Seasonality + Maximum Temperature | 5 | -97.254 | 204.5 | 0.00 | 0.076 |
|  | Isothermality + Maximum Temperature | 4 | -98.314 | 204.6 | 0.12 | 0.071 |
|  | Isothermality + Maximum Temperature + Elevation + Minimum Temperature | 6 | -96.339 | 204.7 | 0.17 | 0.069 |
|  | Isothermality + Maximum Temperature + Precipitation Dry Period | 5 | -97.658 | 205.3 | 0.81 | 0.050 |
|  | Isothermality + Maximum Temperature + Water Deficit Duration | 5 | -97.854 | 205.7 | 1.20 | 0.041 |
|  | Isothermality + Maximum Temperature + Elevation + Minimum Temperature + Water Excess Duration | 7 | -95.871 | 205.7 | 1.23 | 0.041 |
|  | Isothermality + Maximum Temperature + Elevation + Minimum Temperature + Water Excess Severity | 7 | -95.949 | 205.9 | 1.39 | 0.038 |
| Percentage of zoochory species | sesMPD + Minimum Temperature + Water Deficit Severity + Water Excess Duration + Water Excess Severity | 7 | -216.312 | 446.6 | 0.00 | 0.265 |
|  | sesMPD + Water Deficit Severity + Water Excess Duration + Water Excess Severity + Mean Annual Temperature | 7 | -216.324 | 446.6 | 0.02 | 0.262 |
|  | sesMPD + Water Deficit Severity + Water Excess Duration + Water Excess Severity + Maximum Temperature | 7 | -216.589 | 447.2 | 0.55 | 0.201 |
| Percentage of anemochory species | sesMPD + Maximum Temperature + Water Deficit Severity + Water Excess Duration + Water Excess Severity | 7 | -192.657 | 399.3 | 0.00 | 0.114 |
|  | sesMPD + Water Deficit Severity + Water Excess Duration + Water Excess Severity + Mean Annual Temperature | 7 | -192.894 | 399.8 | 0.47 | 0.090 |
|  | sesMPD + Water Deficit Severity + Water Excess Duration + Water Excess Severity + Minimum Temperature | 7 | -192.947 | 399.9 | 0.58 | 0.086 |
|  | sesMPD + Water Excess Duration + Water Excess Severity + Minimum Temperature | 6 | -194.020 | 400.0 | 0.73 | 0.079 |
|  | sesMPD + Water Excess Duration + Water Excess Severity + Minimum Temperature + sesPD | 7 | -193.438 | 100.9 | 1.56 | 0.052 |
|  | sesMPD + Water Excess Severity + Temperature Seasonality + Water Deficit Duration | 6 | -194.622 | 401.2 | 1.93 | 0.044 |
| Percentage of autochory species | sesMPD + Minimum Temperature + Water Deficit Severity + Water Excess Duration + Water Excess Severity | 7 | -149.583 | 313.2 | 0.00 | 0.302 |
|  | sesMPD + Water Deficit Severity + Water Excess Duration + Water Excess Severity + Mean Annual Temperature | 7 | -149.908 | 313.8 | 0.65 | 0.218 |

Table S2. Model-averaged parameter (standard errors) for predicting the ses.PD, ses.MPD, ses.MNTD of forest remnants from the Rio Doce Basin. Significant metrics are highlighted in bold, * is p < 0.05, ** is p < 0.01 and *** is p < 0.001.

|  | ses.PD | ses.MPD | Ses.MNTD |
| --- | --- | --- | --- |
| Intercept | -1.215 ±10.777 | -0.153 ±0.152 | -0.953 ±0.098 |
| PrecAnn | -1.174 ±2.968** | -1.354 ±0.458** | - |
| PrecSeas | 0.062 ±0.477 | - | 0.160 ±0.112 |
| TempMin | 0.233 ±0.656 | -0.049 ±0.161 | -0.110 ±0.321 |
| WaterExcDur | 0.787 ±2.937** | 1.052 ±0.356** | 0.184 ±0.197 |
| WaterExcSev | 0.243 ±0.902 | 0.053 ±0.180 | 0.129 ±0.152 |
| PrecDryP | -0.020 ±0.258 | -0.035 ±0.105 | -0.124 ±0.110 |
| TempAnn | 0.355 ±0.878 | -0.029 ±0.148 | - |
| Isotherm | 0.049 ±0.466 | - | 0.229 ±0.114* |
| WaterDefDur | 0.080 ±0.480 | -0.023 ±0.098 | 0.120 ±0.128 |
| Elevation | -0.020 ±0.193 | -0.069 ±0.317 | 0.454 ±0.337 |
| TempSeas | -0.023 ±0.241 | 0.266 ±0.223 | - |
| TempMax | 0.086 ±0.310 | - | 1.065 ±0.367** |
| WaterDefSev | -0.001 ±0.029 | -0.001 ±0.072 | - |

Table S3. Model-averaged parameter (standard errors) for predicting the ses.PD, ses.MPD, ses.MNTD of forest remnants from the Rio Doce Basin. Significant metrics are highlighted in bold, * is p < 0.05, ** is p < 0.01 and *** is p < 0.001.

|  | Percentual of zoochorous species | Percentual of anemochorous species | Percentual of autochorous species |
| --- | --- | --- | --- |
| Intercept | 72.656 ±0.457 | 20.238 ±0.339 | 7.106 ±0.194 |
| sesMPD | 2.145 ±0.488*** | -1.517 ±0.367*** | -0.6959 ±0.208** |
| sesPD | - | 0.054 ±0.216 | ± |
| sesMNTD | - | - | ± |
| PrecAnn | -1.035 ±1.501 | 0.206 ±0.529 | ± |
| PrecSeas | - | - | ± |
| TempMin | -0.870 ±1.250 | 0.339 ±0.568 | 0.887 ±0.798 |
| WaterExcDur | 2.349 ±0.944* | -1.320 ±0.784 | -0.780 ±0.402 |
| WaterExcSev | -3.625 ±0.832*** | 2.262 ±0.717** | 1.101 ±0.348** |
| PrecDryP | - | - | ± |
| TempAnn | - | - | 0.761 ±0.935 |
| Isotherm | - | - | ± |
| WaterDefDur | - | 0.109 ±0.359 | ± |
| Elevation | - | - | ± |
| TempSeas | - | -0.103 ±0.348 | ± |
| TempMax | -0.851 ±1.491 | 0.316 ±0.681 | ± |
| WaterDefSev | 1.679 ±0.711* | -0.498 ±0.544 | -0.794 ±0.302** |
